# Fast and robust visual object recognition in young children

**DOI:** 10.1101/2024.10.14.618285

**Authors:** Vladislav Ayzenberg, Sukran Bahar Sener, Kylee Novick, Stella F. Lourenco

**Affiliations:** Department of Psychology and Neuroscience, Temple University; Department of Psychology, University of Pennsylvania; Department of Psychology, University of Washington; Department of Psychology, Emory University

**Keywords:** Development, Machine Learning, Object Classification, Shape Perception, Recurrence, Feedback

## Abstract

By adulthood, humans rapidly identify objects from sparse visual displays and across significant disruptions to their appearance. What are the minimal conditions needed to achieve robust recognition abilities and when might these abilities develop? To test this question, we investigated the upper-limits of children’s object recognition abilities. We found that children as young as 3-years-of-age successfully identified objects at speeds of 100 ms (both forward and backward masked) under sparse and disrupted viewing conditions. By contrast, a range computational models implemented with biologically plausible properties or optimized for visual recognition did not reach child-level performance. Models only matched children if they were trained with more data than children are capable of experiencing. These findings highlight the robustness of the human visual system in the absence of extensive experience and identify important developmental constraints for building biologically plausible machines.

## Introduction

Humans extract meaning rapidly from sparse, and often incomplete, visual information. By adulthood, participants identify objects presented as quickly as 100 ms (Grill-Spector & Kanwisher, 2005; Keysers et al., 2001; Thorpe et al., 1996) and they do so across large variations in the visible appearance of objects, such as different orientations (Biederman & Bar, 1999; Biederman & Gerhardstein, 1993) or partial occlusion (Brown & Koch, 2000; Murray et al., 2001). Adult participants also need very little information to infer an object’s identity. For instance, they readily recognize objects when properties such as color, texture, or other internal features are removed— leaving only an object outline (Biederman & Ju, 1988; Elder & Velisavljević, 2009; Wagemans et al., 2008). Moreover, adults maintain high accuracy even when the contours of the object outline are distorted or deleted (Biederman & Cooper, 1991; Wagemans et al., 2012; Wasserstein et al., 1987). Thus, with little visual information the adult visual system is able to quickly and accurately identify objects. What mechanisms support such robust recognition abilities and when do these abilities develop?

Uncovering the mechanisms needed to quickly recognize objects in challenging contexts is difficult when studying only an adult sample. Indeed, when encountering objects in degraded conditions adults may rely on dedicated neural processes that can compensate for missing object information, or they may simply draw on pre-existing knowledge gained from years of visual experience with similar degraded scenarios. For instance, recurrent neural circuits in the ventral visual pathway— the primary pathway underlying visual recognition (DiCarlo et al., 2012)—have been found to support recognition when objects are partially occluded or information is missing (Kok et al., 2016; Lamme & Roelfsema, 2000; Wokke et al., 2013). In this context, the missing portions of an object are perceptually completed via feedback from higher-level areas or lateral input from other portions of the same visual area (Kok et al., 2016; Lamme & Roelfsema, 2000; Wokke et al., 2013). Alternatively, given their breadth of experience, adults may have simply encountered the degraded object in similar conditions previously. Indeed, an extensive perceptual learning literature shows that even minimal exposure to similar objects or contexts improves recognition performance for objects presented in otherwise challenging conditions (Goldstone, 1998; Lu & Dosher, 2022; Squire et al., 2021; Tarr & Bülthoff, 1998). Repeated visual exposure may even alter the response profile of neurons in the ventral pathway, leading to selectivity for commonly encountered objects or stimulus features (Arcaro et al., 2019; Dehaene et al., 2015; Logothetis et al., 1995; Srihasam et al., 2014). In this view, processes such as recurrence may not be needed because observers would have seen similar objects previously and recognition can be accomplished using only a feedforward pass through the ventral pathway.

One approach to help identify the minimal conditions necessary to accomplish object recognition is to study developmental populations (Benton, 2022; Frank, 2023a; Wood, 2013). Specifically, young children provide an ideal subject pool for testing mechanistic questions about object recognition because they have limited visual experience (Clerkin et al., 2017; Huber et al., 2023) and immature visual processing pathways (Ayzenberg et al., 2023; Freud & Behrmann, 2017; Golarai et al., 2007; Nishimura et al., 2015; Scherf et al., 2007). This means that researchers can present stimuli in contexts that children have likely never seen before and track children’s performance in relation to the maturity of their visual system. Together, these factors may allow researchers to isolate the processes that are necessary and sufficient to recognize objects even under challenging conditions. However, little is known about when, and how, such fast and robust recognition abilities develop in childhood.

Yet the use of developmental populations can be challenging for several reasons. One challenge is that children are rarely tested in the same conditions as adults (Frank, 2023a; Tan et al., 2024), making direct comparisons between populations difficult. Specifically, object recognition in human adults, as well as adult non-human primates, is typically tested by presenting objects in a speeded task (100 to 300 ms presentation) and visually masking them (Rajalingham et al., 2018; Wichmann et al., 2017). The development of this standardized approach has been crucial in enabling comparisons between humans and other model organisms (both biological and synthetic), thereby providing an important method by which to explore the mechanisms underlying visual perception (Schrimpf et al., 2020). However, because children’s capacity to follow instructions or stay attentive for long experimental sessions is limited, they are rarely tested in the same challenging conditions as adults, thereby making direct comparisons between organisms difficult and limiting our capacity to use model organisms to explore mechanism. Indeed, the lack of a standardized approach has resulted in unresolved debates regarding the robustness of children’s visual recognition abilities (Nishimura et al., 2009). For instance, research with infants suggests that the processes underlying visual recognition abilities, such as global form perception (Ayzenberg & Lourenco, 2022; Rakison & Butterworth, 1998) and perceptual completion (Johnson, 2004; Valenza et al., 2006), arise early in development (for review, see Ayzenberg & Behrmann, 2024). In contrast, studies with older children suggest that these same processes may not develop until adolescence (Kovács et al., 1999; Scherf et al., 2009). However, because these studies have largely relied on either indirect measures (e.g., looking time) or especially artificial experimental setups (e.g., aligning Gabor wavelets), they are difficult to compare across populations and may not provide a direct or accurate measure of children’s abilities.

In the current study, we sought to determine the upper-bound of young children’s abilities using a challenging object recognition task, similar to the standardized approach typically used with adults (see Figure 1). We did this by requiring 3- to 5-year-olds to identify rapidly presented (100 - 300 ms) 2D outlines of common objects (both forward and backward masked) under different stimulus manipulations. Specifically, we included objects that had complete, undisrupted, contours, as well as objects with perturbed or deleted contours. These stimulus conditions were carefully selected to test the presence of different mechanisms in children. Complete contour objects are those that children of this age have extensive familiarity with via drawings (Long et al., 2024) and likely can identify via a feedforward pass through the ventral pathway (Drewes et al., 2016; von der Heydt, 2015). This condition has the added benefit of providing a baseline for children’s performance in a challenging task. By contrast, perturbed contour objects disrupt the appearance of familiar local visual features and require global form perception, and are challenging for even sophisticated computer vision models (Ayzenberg & Behrmann, 2022a; Baker et al., 2018; Baker et al., 2020). Similarly, deleted contour objects are likely to be unfamiliar to children and they may require perceptual completion as well as recurrent processing within the ventral pathway (Drewes et al., 2016; von der Heydt, 2015). As mentioned previously, these two processes—global form perception and perceptual completion—have historically been thought to develop late in childhood (Davidoff & Roberson, 2002; Jüttner et al., 2013; Kovács, 2000; Kovács et al., 1999; Scherf et al., 2009), but this claim has primarily been tested via indirect methods.

**Figure 1.**
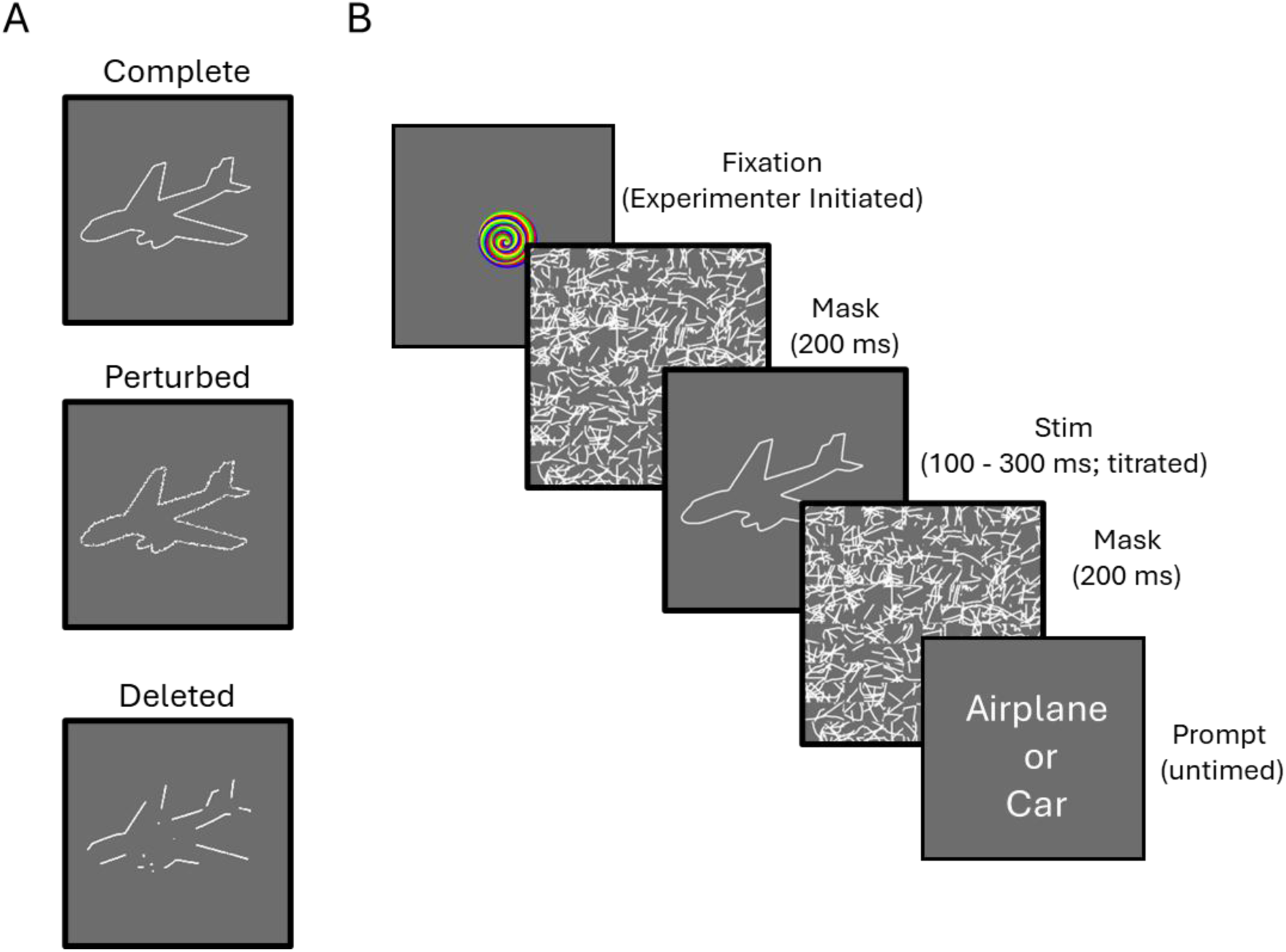
Stimuli and human testing procedure. (A) Children and adults were tested with object outlines that had either complete, perturbed, or deleted contours. (B) On each trial, participants were presented with an object image rapidly (100 – 300 ms duration), which was both forward and backward masked. In the prompt phase, children were asked to verbally indicate which object they saw amongst two possibilities (read by an experimenter). Adults responded by pressing an arrow key that corresponded to each object label.

To further explore what kinds of processes might be necessary to accomplish robust object recognition in childhood, we also compared children to a range of deep neural networks (DNNs; see Table 1). DNNs are computational models that can be trained to accomplish various visual perception tasks. Moreover, they can provide researchers with a well-controlled test bed by which to evaluate theories of visual processing (Doerig et al., 2023; Kriegeskorte, 2015) and their development (Benton, 2024; Pak et al., 2023; Yermolayeva & Rakison, 2014). Indeed, DNNs with recurrent architectures show stronger performance on challenging visual tasks compared to feedforward ones (Linsley et al., 2018; Tang et al., 2018) and they have internal representations that are closely aligned with the primate ventral pathway (Kar et al., 2019; Kietzmann et al., 2019). Other work shows that DNNs trained with naturalistic visual experience and learning objectives are sufficient to recapitulate many human-like visual biases (Geirhos et al., 2018; Sullivan et al., 2020; Vogelsang et al., 2018; M. Vogelsang et al., 2024) and they show human-like performance on a range of object recognition tasks (Mehrer et al., 2021; O’Connell et al., 2023; Sheybani, Hansaria, et al., 2024; Vong et al., 2024). Thus, in the present study, we compared children to a series of DNNs with biologically plausible architectures, visual experiences, or learning objectives, as well as DNNs optimized for performance on various visual perception tasks. By comparing children to DNNs, researchers can better understand the gaps between humans and computational models (Huber et al., 2023; Sheybani, Smith, et al., 2024; Tan et al., 2024; Zaadnoordijk et al., 2020), as well as to help identify models that can be used to inform developmental theory (Cusack et al., 2024; Smith & Slone, 2017; L. Vogelsang et al., 2024; Yuan, 2024).

**Table 1.**
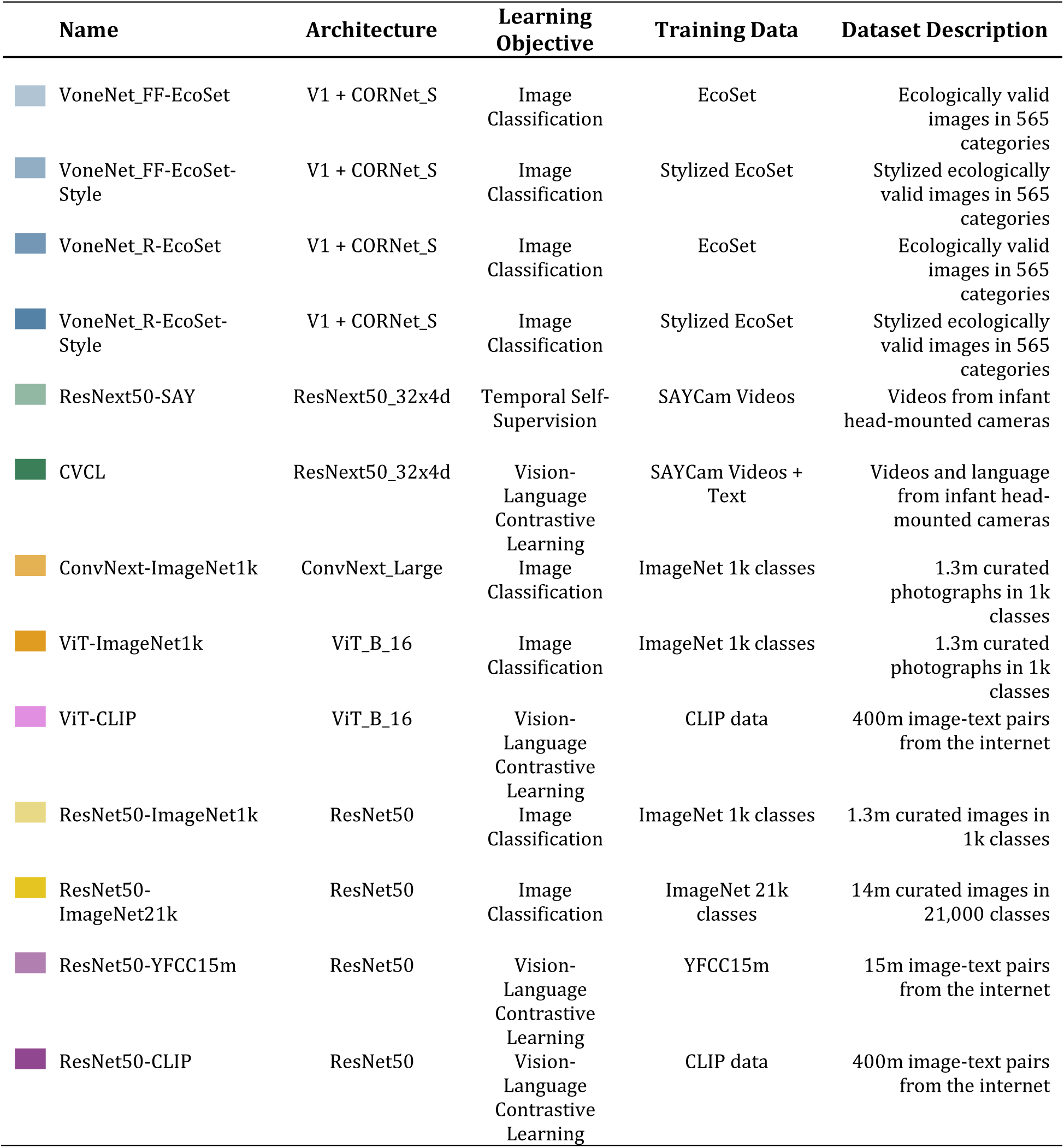
Model description. All models tested in the current study. Colored rectangles correspond to the models in Figure 4.

| Name | Architecture | Learning Objective | Training Data | Dataset Description |
| --- | --- | --- | --- | --- |
| VoneNet_FF-EcoSet | V1 + CORNet_S | Image Classification | EcoSet | Ecologically valid images in 565 categories |
| VoneNet_FF-EcoSet-Style | V1 + CORNet_S | Image Classification | Stylized EcoSet | Stylized ecologically valid images in 565 categories |
| VoneNet_R-EcoSet | V1 + CORNet_S | Image Classification | EcoSet | Ecologically valid images in 565 categories |
| VoneNet_R-EcoSet-Style | V1 + CORNet_S | Image Classification | Stylized EcoSet | Stylized ecologically valid images in 565 categories |
| ResNext50-SAY | ResNext50_32x4d | Temporal Self-Supervision | SAYCam Videos | Videos from infant head-mounted cameras |
| CVCL | ResNext50_32x4d | Vision-Language Contrastive Learning | SAYCam Videos + Text | Videos and language from infant head-mounted cameras |
| ConvNext-ImageNet1k | ConvNext_Large | Image Classification | ImageNet 1k classes | 1.3m curated photographs in 1k classes |
| ViT-ImageNet1k | ViT_B_16 | Image Classification | ImageNet 1k classes | 1.3m curated photographs in 1k classes |
| ViT-CLIP | ViT_B_16 | Vision-Language Contrastive Learning | CLIP data | 400m image-text pairs from the internet |
| ResNet50-ImageNet1k | ResNet50 | Image Classification | ImageNet 1k classes | 1.3m curated images in 1k classes |
| ResNet50-ImageNet21k | ResNet50 | Image Classification | ImageNet 21k classes | 14m curated images in 21,000 classes |
| ResNet50-YFCC15m | ResNet50 | Vision-Language Contrastive Learning | YFCC15m | 15m image-text pairs from the internet |
| ResNet50-CLIP | ResNet50 | Vision-Language Contrastive Learning | CLIP data | 400m image-text pairs from the internet |

## Results

### Child performance

What are the upper limits of children’s visual recognition abilities? We tested this question by presenting 3-, 4-, and 5-year-olds with a challenging recognition task where they were required to identify rapidly presented objects (100 to 300 ms) that were both forward and backward masked (see Figure 1 and Supplemental Video). Moreover, objects could be presented with either complete, perturbed, or deleted contours.

Overall, we found that children’s overall performance was above chance (0.5) in all conditions, even when stimuli were presented at the fastest speed of 100 ms (*ps* < .001, *ds* > 0.65; see Figure 2), suggesting that object recognition is fast and robust from a young age.

**Figure 2.**
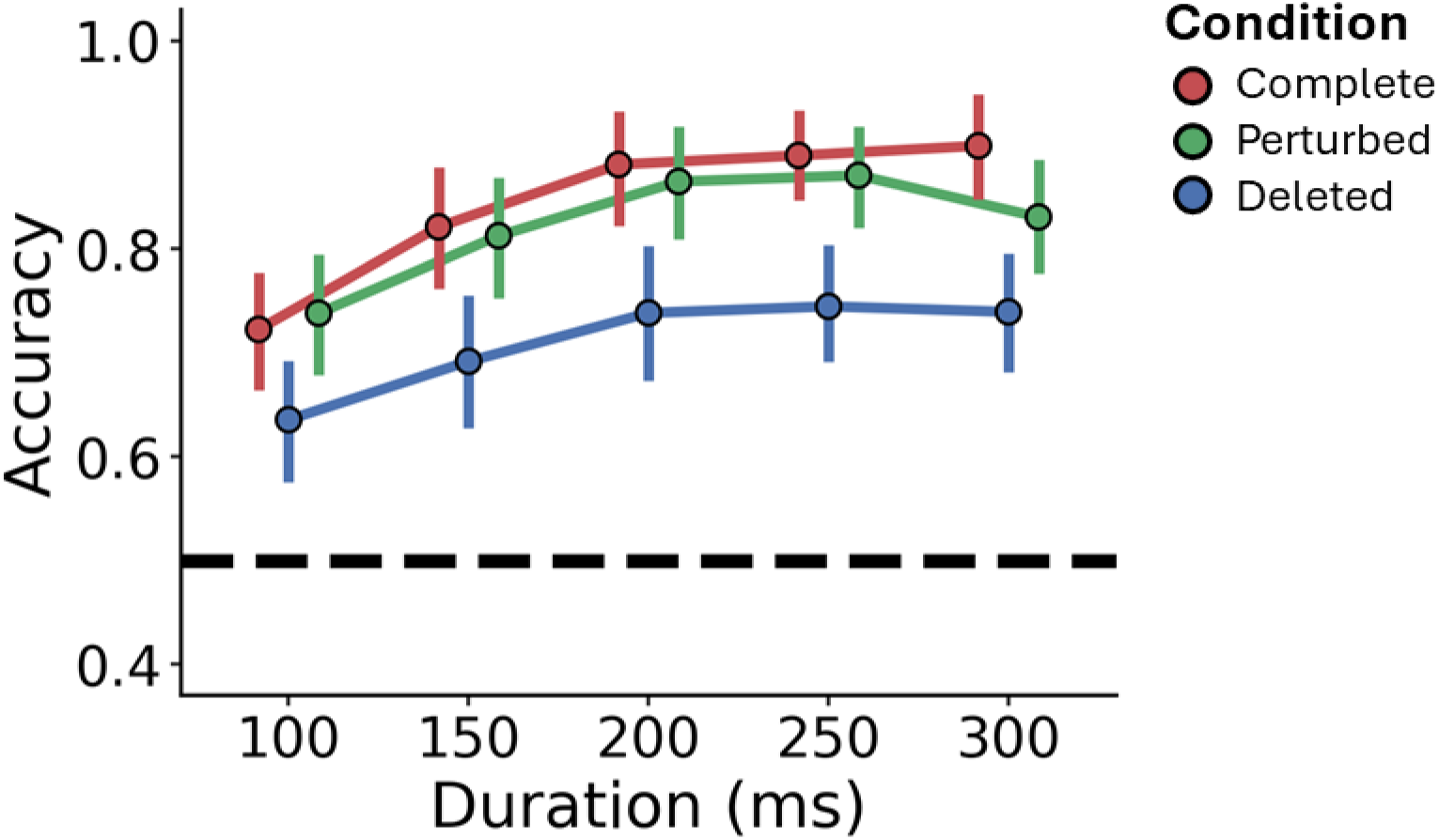
Children’s performance in each condition. Across age, children performed above chance for each condition at each duration. Error bars indicate standard error of the mean (SEM). Dotted black line indicates chance performance (0.5).

Next, we examined whether there were differences in performance as a function of stimulus condition (complete, perturbed, or deleted), presentation time (durations: 100-300 ms), and child’s age. A repeated measures ANCOVA, with age as a covariate, revealed main effects of condition, *F*(2, 122) = 16.62, *p* < .001, η_p_^2^ = .21, duration, *F*(4, 488) = 2.64, *p* = .033, η_p_^2^ = .02, and age, *F*(4, 488) = 28.25, *p* < .001, η_p_^2^ = .19. There were no significant interactions between factors (*ps* > .314). Overall, participants’ performance was higher at slower durations than faster ones. Moreover, post-hoc comparisons (Holm-Bonferroni corrected) revealed that performance was overall worse for the deleted condition than either complete or perturbed conditions (*ps* < .001, *ds* > 0.66; see Figure 2). There was no difference between complete and perturbed conditions (*p* = .502).

When age is converted into a categorical variable (3, 4, and 5 years), we found that 3-year-olds performed worse than 4- and 5-year-olds (*ps* <.001; *ds* > 0.70), but there was no difference between 4- and 5-year-olds’ performance (*p* = .136, *d* = 0.20; see Figure 3). An additional comparison to adults revealed that adults performed better than children of every age and condition (*ps* < .001; *ds* > 0.78; see Figure 3).

**Figure 3.**
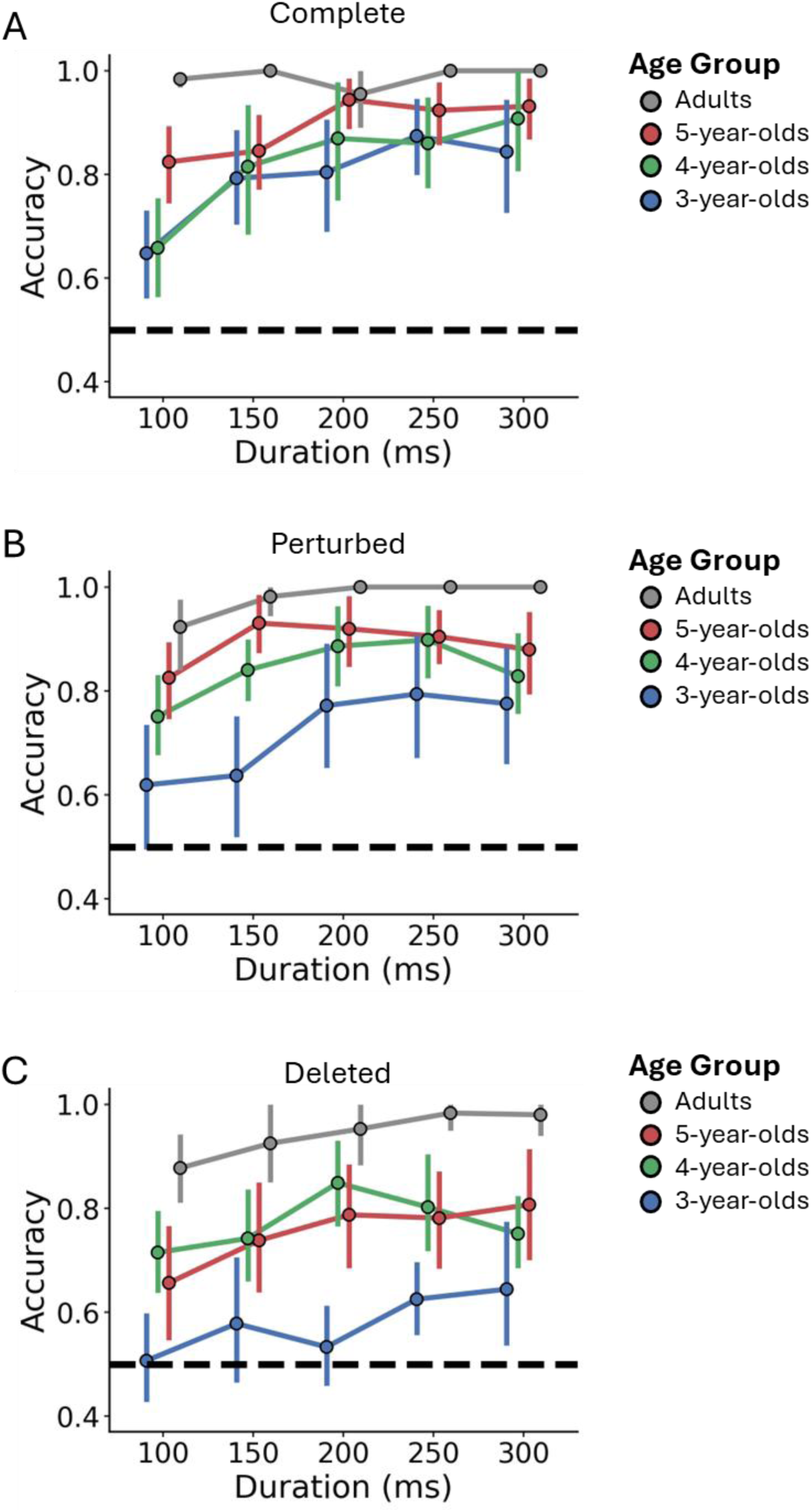
Performance in each condition by age group. (A) In the complete condition, participants of all ages performed above chance, even at the fastest speeds. (B) In the perturbed condition, 4- and 5-year-olds performed above chance at all speeds, whereas 3-year-olds were only above chance when durations were 200 ms and slower. (C) In the deleted condition, 4- and 5-year-olds performed above chance at all speeds, whereas 3-year-olds only performed above chance at the slowest speeds (250 ms and 300 ms). Error bars indicate standard error of the mean (SEM). Dotted black line indicates chance performance (0.5).

To provide a more detailed understanding of the development of object recognition, we examined performance separately for each age group and condition (see Figure 3). We found that, like adults, 4- and 5-year-olds performed above chance in every condition and at all durations (*ps* < .038, *ds* > 0.63). By contrast, 3-year-olds performed above chance at all durations of the complete condition (*ps* < .007, *ds* > 1.00), but only at speeds of 200 ms or slower in the perturbed condition (*ps* < .001, *ds* > 1.27) and 250 ms or slower in the deleted condition (*ps* < .007, *ds* > 0.98). That children, especially in the youngest age group, performed worse in the deleted condition may suggest that perceptual completion and recurrent processing in the ventral pathway may still be immature in preschool-aged children.

Altogether, these findings suggest that by 4 years of age, children rapidly extract meaning from sparse visual displays, including when there is missing information. That said, even 3-year-old children performed above chance in most conditions, despite stimuli being presented as fast as 100 ms with both forward and backward masking. Nevertheless, children’s object recognition abilities are not fully mature, as even 5-year-old children performed worse than adults.

### Model comparison

How does children’s performance compare to biologically plausible and performance-optimized DNNs? To answer this question, we compared children (and adults) to DNN models with different types of architectures (e.g., feedforward vs. recurrent), visual experience (e.g., curated vs. variable visual experience), and learning objectives (e.g., classification vs. vision-language alignment), as well as DNNs optimized for object classification tasks (see Table 1). Participants’ performance in each age group (3, 4, 5 years, and adult) was split into fast (mean of 100 and 150 ms durations) and slow (mean of 200 and 250 ms durations) durations.

Overall, models performed above chance in every condition, regardless of their architecture, training experience, or learning objective (see Figure 4; Supplemental Tables 2-4). Like humans, these models generally performed best on the complete condition, and worst on the deleted condition (see Supplemental Figure 2). However, whereas humans did not show a significant difference between complete and perturbed conditions, models generally performed worse on the perturbed condition. Amongst biologically plausible models, recurrent models (VoneNet_R) generally outperformed feedforward models (VoneNet_FF), which is consistent with the hypothesis that recurrence may be crucial for robust object recognition. Training with variable visual experience (i.e., Stylized EcoSet) showed a benefit, but only in the deleted condition. Interestingly, VoneNet_R performance was as strong, or stronger, than that of ConvNext-ImageNet1k and ViT-ImageNet1k, even though VoneNet models have fewer parameters (VoneNet_R: 55m params, ConvNext: 198m params; ViT: 86m params). This suggests that implementing biologically plausible properties into computational models may lead to improvements in the performance and efficiency of DNNs. There were not major improvements, however, from models trained with naturalistic videos from infant head-mounted cameras, regardless of whether they were trained purely on videos (ResNext50_SAY) or when there was a vision-language objective (CVCL).

**Figure 4.**
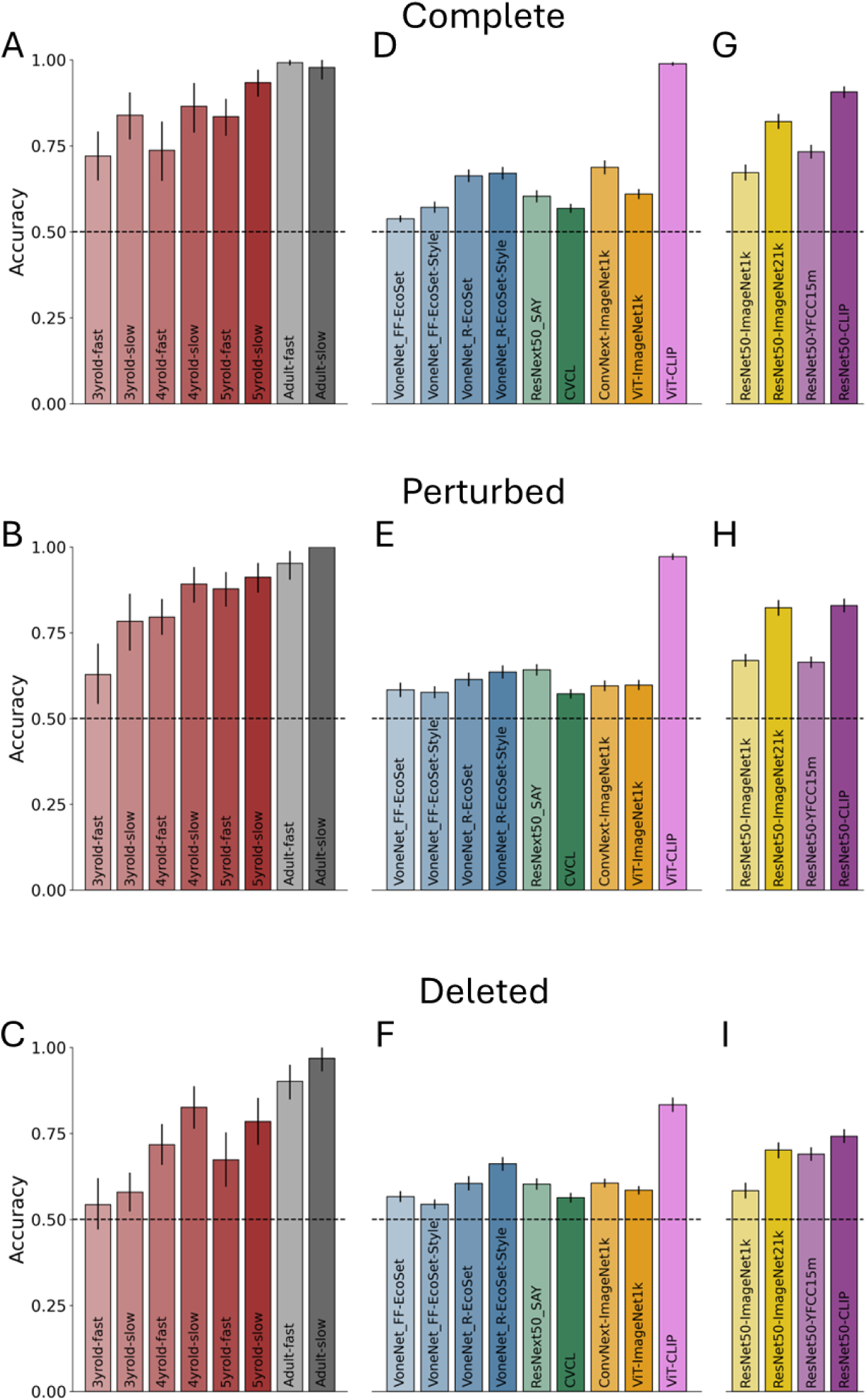
Performance of humans and models in the (top) complete, (middle) perturbed, and (bottom) deleted contour conditions. (A-C) Human data for each age (red: children; gray: adults) were aggregated into fast (100 ms & 150 ms) and slow (200 ms & 250 ms) stimulus durations. Humans were compared to (D-F) biologically plausible (blue: ventral-like architecture; green: trained on infant experience) and performance-optimized (orange: classification objective; violet: vision-language objective) models, as well as (G-I) models selected to disambiguate between the contributions of training scale and a vision-language objective (yellow: classification objective; purple: vision-language objective). The Y-axis indicates classification accuracy. The dotted line indicates chance performance (0.5) Error bars depict 95% confidence intervals.

However, we found that neither biologically plausible, nor performance-optimized classification models, consistently matched the performance of children across all conditions (see Figure 4). The only instance where one of these models outperformed a child was VoneNet-R_ EcoSet-Style, which outperformed 3-year-olds, but only when they were tested in the fastest duration. The one model whose performance did consistently match, or surpass, children was ViT-CLIP, a vision-language model trained to associate images and text via a contrastive learning objective. Indeed, ViT-CLIP performance matched that of adults in complete and perturbed conditions (but not deleted), and surpassed all children for the complete and perturbed conditions, and was comparable to 4- and 5-year-olds in the deleted conditions for the slow durations.

### Language or Scale?

What properties of ViT-CLIP account for its strong visual recognition performance? One possibility is that a learning objective which pairs linguistic information with images, results in a more robust visual representation. Indeed, classic developmental work has suggested that language bootstraps object recognition by guiding children’s attention to diagnostic object properties (Smith et al., 1996; Smith et al., 2002). However, the poor performance of CVCL, a model trained to associate naturalistic video with linguistic information from infants, suggests that a vision-language objective on its own may not be sufficient for a DNNs.

An alternative possibility is that the success of ViT-CLIP is attributed to its extensive visual experience. Indeed, in contrast to the classification models tested above, which were trained with ImageNet (1.3 m images) or EcoSet (1.5 m images), CLIP was trained with a proprietary dataset of over 400 m images from the internet (see Table 1). For comparison, if, from the moment of birth, a child were to see a new object image every second of their life for six consecutive years, they would have seen only 189 m images--less than half the number of images that CLIP was trained on. This back-of-the-envelope estimate ignores the fact that infants, conservatively, sleep an average of 14+ hours per day in the first year of life and 3- to 5-year-olds sleep at least 10 hours per day (*Centers for Disease Control and Prevention*, 2024; Paruthi et al., 2016).

To test whether data size or a vision-language learning objective contributed to model performance, we tested four additional models. Specifically, we tested ResNet50 models that were pre-trained on image datasets of similar size (ImageNet21k: 14 m images; YFFC15m: 15 m images), with one using a standard object classification objective and dataset (ImageNet21k) and the other using a vision-language objective to align image and text embeddings (YFFC15m). As baselines, we also included a ResNet50 model trained to classify images from the standard ImageNet1k dataset (1.3 m images) and one trained with the same procedure and dataset as CLIP (ResNet50-CLIP; 400 m images).

Consistent with the prior analysis, we found that ResNet50-CLIP showed the strongest performance compared to the other models, and it matched (or surpassed) children in almost every condition (see Figure 4G-I). When comparing ResNet50-CLIP to other models, our results suggest that human-level performance is related to training size, not a vision-language learning objective. Indeed, the performance of a ResNet50 model trained on ImageNet21k, showed a large improvement over a base ResNet50 model trained on ImageNet1k. Moreover, ResNet50-ImageNet21k outperformed ResNet50-YFFC15m on the complete and perturbed image conditions, and its performance often matched children when tested at the fastest speeds. Taken together with the previous model comparisons, these findings suggest that the strongest driver of DNN performance may be the scale of the training data, rather than different biologically plausible architectures, naturalistic training data, or learning objectives.

### General Discussion

In the current study, we sought to understand the development of robust object recognition in young children. Our results demonstrate that young children succeed at identifying objects from sparse visual displays, at speeds as fast as 100 ms and even when the contours are disrupted. Direct comparisons to a range of computer vision models revealed that children outperformed both biologically plausible and performance-optimized DNNs. Only by exponentially increasing a model’s training data, beyond what children can realistically experience, did models surpass children’s performance. Altogether, these findings suggest that object recognition is fast and robust from early in development, but gaps remain in our ability to approximate these processes with current computational models.

### Visual recognition in young children

We found that by 4 years of age, children identified objects whose local visual features were disrupted via contour perturbations or deletions, even when presented as rapidly as 100 ms, and even though stimuli were forward and backward masked. Remarkably, even 3-year-old children performed above chance, though they required somewhat slower presentation times when object contours were disrupted. These findings suggest that young children exhibit robust recognition in which they readily extract global form across variations in local features.

Our results stand in stark contrast to a collection of studies in which the findings suggested a protracted development of object recognition abilities (Nishimura et al., 2009). Specifically, these studies found that the ability to ignore local features, so as to represent global form, or to accomplish perceptual completion across disconnected contours remained difficult for children as old as 10 years of age (Davidoff & Roberson, 2002; Jüttner et al., 2013; Kovács, 2000; Kovács et al., 1999; Scherf et al., 2009). These findings have been mixed, however. Other studies with similar methods and age groups found the opposite, that children prioritized global form over local features (Wakui et al., 2013).

Instead, our findings are consistent with a rich literature with younger children that suggests object recognition abilities that develop earlier in life. For instance, 2-year-olds preferentially use shape information to categorize objects across changes in color and texture information (Abdelrahim & Frank, 2024; Landau et al., 1988), and with linguistic scaffolding young children generalize object identities to sparse caricature versions of common objects (Pereira & Smith, 2009; Smith, 2003, 2009). Furthermore, research with infants has shown that by 3 months of age, they identify novel objects from orientations not previously seen (Kraebel & Gerhardstein, 2006; Mash et al., 2007; Ruff, 1978; Slater et al., 1991; Slater & Morison, 1987; Slater et al., 1984), and older infants are capable of categorizing novel objects across variations in contours (Ayzenberg & Lourenco, 2022; Mareschal & Quinn, 2001; Quinn, Slater, et al., 2001; Turati et al., 2003). Indeed, by the end of the first year of life, children can infer an object’s category from only its shape silhouette (Quinn, Eimas, et al., 2001), suggesting that they do not rely on color, texture, or internal local features to recognize objects. Finally, other work has shown that infants are capable of inferring the complete shape of an object even when it is partially occluded (Johnson & Aslin, 1995; Kellman & Spelke, 1983; Kellman et al., 1998; Slater et al., 1990). However, studies with infants cannot measure speed of processing and rely on indirect looking time measures to infer processes like recognition and are difficult to compare to different populations. Our work shows that by at least preschool age, children are capable of identifying rapidly presented objects from sparse and disrupted visual displays.

We would not suggest, however, that children’s visual recognition abilities are fully mature at preschool age. Indeed, we generally found that children’s performance in all conditions and at all trial durations, improved with age, with all children performing worse than adults. The effect of age was most pronounced in the deleted condition. Specifically, in the deleted condition, 3-year-olds performed at chance at most speeds, whereas by 4 years of age, children performed above chance at all speeds. Given that the recognition of contour-deleted objects may require perceptual completion and recurrent processing, the developmental change observed here may indicate underdeveloped recurrent connections in 3-year-old children. Thus, one interpretation of our results, which is consistent with the existing literature from both infants and older children, is that children have the capacity for robust object recognition from early in development, but this ability does not become fully adult-like until adolescence (Kovács, 2000).

### Using children as a benchmark

A major goal of the current study was to compare children to computational models to explore what mechanisms might support the development of visual recognition abilities and to assess the gaps between humans and machines. Such comparisons are crucial if DNNs are to be used as tools to explore the cognitive and neural mechanisms underlying visual recognition in humans (Bowers et al., 2023; Doerig et al., 2023; Schrimpf et al., 2020). Although we found that factors like a biologically plausible recurrent architecture (VoneNet-R) allowed models to match, or even outperform much larger DNNs optimized for image classification (e.g., ConvNext), these models rarely matched child performance. Instead, the only models to consistently match, or surpass, children’s performance were those trained on especially large datasets (e.g., ImageNet21k, CLIP).

Could children’s performance in the current study reflect the scale of their visual experience? Our analyses, as well as existing research with children, suggest that this possibility is unlikely. First, as described in the results, even a conservative estimate of children’s object exposure reveals that they are unable to receive as much visual experience as a CLIP model until they are well into adolescence. Moreover, studies analyzing children’s visual experience from head-mounted cameras show that children are exposed to relatively few distinct objects. Indeed, in the first year, just 10 common objects occupies approximately 1/3 of their visual experience (Clerkin et al., 2017), with another large portion accounted for by three faces (Jayaraman et al., 2015; Jayaraman & Smith, 2019), hands (Long et al., 2022), and visually simple architectural features of a scene (e.g., bright ceiling light; Anderson et al., 2024). Instead, children’s experience with these few objects is extensive, such that they densely sample many views of the same few objects (James et al., 2014; Smith et al., 2018). Even by 5 years of age, children likely have less object knowledge than a smaller model like ResNet50-ImageNet21k. Indeed, children’s total vocabulary at this age is approximately 10k words (Shipley & McAfee, 2023), less than half of the number of object classes in ImageNet21k. These findings contribute to a growing literature that illustrates how children develop sophisticated perceptual (Ayzenberg & Lourenco, 2022; Huber et al., 2023; Sheybani, Smith, et al., 2024), linguistic (Frank, 2023b; Tan et al., 2024), and other high-level cognitive abilities (Cherian et al., 2023; Goddu & Gopnik, 2024; Yiu et al., 2024) from far less data than state-of-the-art machine learning models currently require (Zador, 2019).

What, then, might account for the discrepancy between children and DNNs? One possibility is that we only tested recurrent models that implement local, within-layer, recurrence where information is processed by the same layer multiple times. By contrast, recurrent processing in the human ventral pathway occurs both within and between cortical areas, a process also known as top-down feedback (Bar et al., 2006; Lee et al., 1998). Indeed, top-down feedback may be particularly important for perceptual completion because it exploits the larger-receptive fields of higher-level visual areas (Lee et al., 1998; von der Heydt, 2015; Wokke et al., 2013). Moreover, it provides a mechanism by which prior knowledge from areas such as the pre-frontal cortex can be used to make predictions about the content of ambiguous displays (Bar, 2004; Hardstone et al., 2021; Teufel et al., 2018). Thus, top-down feedback offers both a perceptual mechanism by which to fill-in missing information and a process by which to efficiently apply prior experience to the current scenario.

Another possibility is that the current DNNs only approximate the ventral visual pathway, which, on its own, may not be sufficient for robust object recognition. Specifically, accumulating evidence suggests that the ventral pathway, like DNNs, is most sensitive to local visual features (Ayzenberg & Behrmann, 2022a; Jagadeesh & Gardner, 2022; Long et al., 2018; Wang & Ponce, 2022), and it has difficulty representing a complete object form without input from the dorsal visual pathway and prefrontal cortex (Ayzenberg et al., 2022; Bar et al., 2006; Freud et al., 2016; Kar & DiCarlo, 2021; Romei et al., 2011; Rose & Ponce, 2024; Zachariou et al., 2017). Dorsal visual areas develop early (Ayzenberg et al., 2023; Ayzenberg et al., 2024; Bourne & Rosa, 2006) and support the formation of global shape percepts by representing the spatial arrangement, rather than the appearance, of object parts (Ayzenberg & Behrmann, 2022b; Romei et al., 2011). And, as mentioned previously, pre-frontal areas may be crucial for applying prior knowledge to solve visual recognition tasks (Bar et al., 2006), though these areas may develop later (Kolk & Rakic, 2022; but, see Ellis et al., 2021). Nevertheless, models that implement the processes of dorsal and pre-frontal cortices may better approximate human recognition abilities across development.

A final possibility is that the *kind* of visual experience children receive, rather than the quantity, is crucial for robust object recognition. As mentioned previously, children are exposed to relatively few objects in the first year of life, but their experience with these objects is far more variable and extensive (Smith et al., 2018). Indeed, DNNs trained on children’s views of objects from head-cam videos perform better than those trained on the same objects using adult viewpoints (Bambach et al., 2017), suggesting that child experience provides better training data than that of adults. Moreover, other work shows that building in constraints of the newborn visual system, such as infants’ initially blurry and low-saturation vision (Vogelsang et al., 2018; M. Vogelsang et al., 2024), improves the robustness of DNNs. Although our data suggest that naturalistic visual experience alone is insufficient for DNNs to match child performance, integrating both biologically plausible architectures and naturalistic experiences may help close the gaps between humans and machines.

### Conclusion

Whereas extensive research has explored the visual capacities of human adults, much less is known about the development of such capacities in early childhood. A developmental perspective is crucial for understanding the organization of the adult visual system because it describes the necessary conditions that lead to robust visual recognition and may even provide a blueprint by which researchers can train more human-like machines. In the current study, we found that young children already exhibit remarkable object recognition abilities, identifying objects quickly and across various contexts. By contrast, a range of biologically plausible or performance-optimized DNN models had difficulty identifying objects in the same conditions. These findings illustrate that mechanisms underlying robust visual recognition remain poorly understood. Yet, by taking a developmental perspective, we may ultimately uncover the processes by which humans attain sophisticated perceptual abilities.

## Methods

### Participants

For human children, sample sizes and testing procedures were preregistered (https://aspredicted.org/Z6B_PF4) on the basis of pilot testing. We sought to test 135 children ages 3 through 5 years of age. Five additional children were included because they were scheduled before the target sample size was met. An additional 12 children were excluded from the analyses for not following instructions (*n* = 3), failing to complete the task (*n* = 5), technical issues (*n* = 3), or because of developmental disability (*n* = 1). The five additional children were thus retained to compensate for those who were excluded. In total, data from 128 children (*M_age_* = 4.62 years, range = 3.05 – 5.95; 64 girls and 64 boys) were analyzed. Children were randomly assigned to one of three stimulus conditions (between-subjects: complete, perturbed, or deleted), with an equal number of children (*n* = 42) per condition. For comparison, we also recruited 30 adult participants (10 per condition) online. One participant did not finish the task and thus 29 participants were analyzed. Age and sex were not collected for adults. All participants were tested online through Children Helping Science (children) or Prolific (adults).

### Models

For computational models, we selected a combination of models based on biologically plausible properties or performance on visual recognition tasks. See Table 1 for all models.

To examine the possible contributions of feedforward and recurrent neural circuits, we implemented a shallow feedforward and recurrent VoneNet architecture (Dapello et al., 2020). VoneNet is a convolutional DNN whose architecture is based on the primate ventral visual pathway, with distinct layers corresponding to V1, V2, V4, and IT. What further distinguishes VoneNet from other DNNs is that layer V1 directly simulates the organization of primary visual cortex by implementing a fixed weight front-end comprised of a Gabor filter bank. The inclusion of a biologically plausible front-end was shown to improve the adversarial robustness of DNNs, as well as VoneNet’s match to monkey ventral pathway (Dapello et al., 2020). Feedforward or recurrent processing was implemented into each VoneNet architecture by manipulating the number of times a stimulus was passed through each layer on a single presentation (i.e., recurrent steps). In the feedforward architecture, each stimulus was processed only once by each layer. For the recurrent architecture, the number of recurrent steps for each layer (V2: 2, V4: 4, IT: 2) was selected on the basis of prior work (Dapello et al., 2020).

In addition, to explore the contributions of different kinds of visual experience, each VoneNet architecture was trained on either EcoSet, an ecologically-valid stimulus set comprised of 565 basic-level categories (Mehrer et al., 2021), or stylized EcoSet, which uses style transfer techniques to randomly vary the color and texture of each EcoSet image (Gatys et al., 2016). We specifically included stylized EcoSet because prior work found that conventional DNNs are biased to use texture information, but training on style transfer images induces a more human-like shape bias in the models (Geirhos et al., 2018). All models were trained in a supervised manner to classify images into 565 categories. Models were trained in PyTorch (Paszke et al., 2019) using the procedure described in Dapello et al. (2020). Thus, four biologically plausible models were trained for this study: feedforward models trained either on EcoSet or Stylized-EcoSet, and recurrent models trained on the same two datasets. The use of these models allowed us to explore two possible mechanisms underlying visual recognition: recurrent processing and variable visual experience.

We also tested two other models with biologically plausible properties. These included a ResNext50 model trained on the SAYCam dataset (Sullivan et al., 2020), a dataset consisting of naturalistic head-cam videos from three infants (infant: ‘S’, ‘A’, and ‘Y’), using temporal self-supervision method (from here on referred to as ResNext50-SAY). Furthermore, to examine whether language may bootstrap visual learning, we tested the CVCL (Child’s View for Contrastive Learning) model, a contrastive ResNext50 model that was trained to associate linguistic-utterances with co-occurring visual information from the head-cam videos of a single infant from the SAYCam dataset (Vong et al., 2024). CVCL was included because it successfully uses biologically plausible training data and learning objective to recapitulate the visual abilities of a young child.

To estimate the gaps (or lack thereof) between child and machine perception, we also tested several performance-optimized object classification models (as of March 2024). Specifically, we tested a Vision Transformer (ViT), which was trained on ImageNet using a transformer architecture (Dosovitskiy et al., 2020), ConvNext, which is a state-of-the-art convolutional DNN that was optimized using the best-practices from the last decade of machine learning to maximize visual recognition performance on ImageNet (Liu et al., 2022), and CLIP, a vision-language model using the ViT architecture that learned to associate 400m images and text pairs using a contrastive learning objective (Radford et al., 2021).

Finally, to examine the role of language and training size on performance, we also tested four versions of ResNet50, trained to either do image classification (He et al., 2016; Ridnik et al., 2021) or text-image association (Cherti et al., 2023; Radford et al., 2021). Specifically, we tested a ResNet50 trained to classify images from standard ImageNet (1000 classes, 1.4m images) or ImageNet 21k (21k classes, 14m images), as well as a ResNet50 architectures trained to associate text-image pairs from YFFC 15M (15m images) or the CLIP training set (400m images).

### Stimuli

Thirty unique objects (15 animate; 15 inanimate) were selected for the object recognition task created for this study. Objects were selected from the Snodgrass and Vanderwart (1980) image set by sampling age-appropriate nouns from the Peabody Picture Vocabulary Test (PPVT-4; Dunn & Dunn, 2007). See Supplemental Table 1 for the list of objects used. Each object image was transformed into an outline and adapted into three stimulus conditions: complete, perturbed, and deleted contours. Stimuli with complete contours were adopted as is from Snodgrass and Vanderwart (1980). Perturbed stimuli were created by applying the ripple distortion (amount = 200%; size = medium) to the complete contour stimuli in using image editing software (Photopea; https://www.photopea.com/). The deleted contour stimuli were created by removing 50% of the contours from each complete stimulus using salience scores based on the medial axis of the object (Rezanejad et al., 2019). To minimize visual discomfort during the testing session, stimulus contours were presented as white lines on a gray background (see Figure 1).

### Human testing procedure

Each child was tested individually by an experimenter over Zoom using a two-alternative forced-choice (2AFC) procedure. Trials began with a colorful fixation stimulus, which remained onscreen until children attended to it (as determined by the experimenter). The stimulus display was sandwiched by forward and backward masks; the response prompt followed (Figure 1B). Masks were created by randomly overlaying object outlines across the image frame and then box scrambling the result.

During presentation of the response display, the experimenter asked the child whether they saw the target stimulus or a distractor (e.g., “Did you see an airplane or a car?”; order randomized across trials). The distractor was always another object from the stimulus set with the same animacy (e.g., animate target paired with animate distractor; randomly selected), ensuring that children could not rely on low-level stimulus features to identify the objects. The response phase was followed by feedback indicating whether the child responded correctly (green check) or incorrectly (gray square).

Duration of stimulus displays varied between 100-300 ms. Durations were determined via a titrated procedure. At the beginning of each session, objects were presented for 300 ms. If children correctly identified objects on three consecutive trials, then the duration decreased by 50 ms (capped at 100 ms). However, if children incorrectly identified objects for three consecutive trials, then the stimulus duration increased by 50 ms (capped at 300 ms).

Prior to testing, children were introduced to the task in a practice phase where they were required to identify slowly presented (1 s) photographs (e.g., shoes) using the same instructions as above.

Adults were tested using an identical procedure, except that testing was conducted asynchronously without an experimenter present. Adults made their response by pressing the arrow key that corresponded to either the target or distractor label.

### Model testing procedures

We tested model recognition accuracy using the same stimulus set presented to children and with a comparable 2AFC procedure. In the object recognition task, children were required to determine which of two semantic labels corresponded to the target stimulus (e.g., “airplane” or “car”). To accomplish this task, children must be able to match the stored object representation that is associated with each label (e.g., examples of previously seen airplanes) with the stimulus image they saw during the trial.

To approximate this process, we trained machine learning classifiers using the feature activations from the penultimate layer of each model on naturalistic images of each object and then tested them on each stimulus display in a pairwise fashion (e.g., train on photographs of airplanes and cars; test on the perturbed airplane stimulus). For each stimulus category, we provided 500 images for training (15,000 images total). Training images were randomly selected from the EcoSet (Mehrer et al., 2021), ImageNet21k (Ridnik et al., 2021), and Open Images (Kuznetsova et al., 2020) data sets. Visual inspection of the images showed that they comprised both photographic (i.e., real) and stylized (e.g., cartoons) examples, making them comparable to a child’s viewing experience.

Because there can be significant differences in the performance of a model based on which classifier is used or how many examples are used to train the classifier, we tested each model using six common classifiers (support vector machine, logistic regression, ridge regression, naïve bayes, K-nearest neighbors, and nearest centroid). We specifically included the K-nearest neighbors and nearest centroid classifier because these methods may also approximate the use of exemplar and prototype representations in humans, respectively. Furthermore, we parametrically varied the number of images used for classifier training (5, 10, 25, 50, 100, 150, 200, 250, and 300). For each stimulus, training and testing was done using a 20-fold cross validation regime (stratified k-fold). For each model, final comparisons with children were made by selecting the best performing version of the model across all classifiers and quantity of training.

### Statistical Analyses

#### Preprocessing of child data

In the current task, we used a titrated procedure to determine the fastest speed at which children could identify images. However, the use of a titrated procedure meant that not all children contributed trials at every stimulus duration. Indeed, survival analyses revealed that 3-year-olds had a lower probability of reaching the fastest speeds in the perturbed and deleted conditions than 4- and 5-year-olds (but not the completed condition; see Supplemental Figure 1). However, if we only included data from children that reached each speed, then this would likely overestimate children’s performance (particularly at the fastest speeds) because it would exclude every participant that might otherwise perform at chance. To address this issue, we imputed data for each missing trial using a conservative estimate that assumes children’s performance for those trials would be below chance. This value was computed for participants with missing data by subtracting the standard error of the mean of their performance on existing trials from 0.5. This strongly penalizes the overall group estimate of performance for those trials and ensures that we are not overestimating the strength of children’s performance.

#### Model comparison

Humans and models were compared on the basis of overlapping confidence intervals (95% CIs). Human data were split into four age groups (3-, 4-, 5-year-olds, and adults) and two speeds (fast: mean accuracies at 100 ms and 150 ms; slow: mean accuracies at 200 ms and 250 ms). Performance for models was computed as mean decoding accuracy across all pairwise comparisons of objects from the best performing version of the model across all classifiers and quantity of training examples. Confidence intervals (CIs) for humans and models were computed using a bootstrapping procedure (10,000 resamples).

## Conflicts of interest

The authors declare no conflicts of interest

## Notes

### Competing Interest Statement

The authors have declared no competing interest.

### Summary of Updates

We were required to replace the current version of the manuscript with an older version to meet the publication requirements of a journal.

