## Supplemental Materials for "Fast and robust visual object recognition in young children"

**Supplemental Table 1.** List of objects tested in the current study

| Object | Animacy Type |
|---|---|
| Airplane | Inanimate |
| Apple | Inanimate |
| Bear | Animate |
| Bicycle | Inanimate |
| Bird | Animate |
| Butterfly | Animate |
| Cat | Animate |
| Chair | Inanimate |
| Cow | Animate |
| Dog | Animate |
| Duck | Animate |
| Elephant | Animate |
| Fish | Animate |
| Flower | Inanimate |
| Foot | Animate |
| Fork | Inanimate |
| Frog | Animate |
| Hand | Animate |
| Guitar | Inanimate |
| Key | Inanimate |
| Kite | Inanimate |
| Lamp | Inanimate |
| Leaf | Inanimate |
| Rabbit | Animate |
| Spoon | Inanimate |
| Squirrel | Animate |
| Tree | Inanimate |
| Truck | Inanimate |
| Turtle | Animate |
| Umbrella | Inanimate |

**Supplementary Table 2.** Human and model accuracy on the complete condition. CI Lower and Upper refer to upper and lower bounds of the 95% confidence intervals. Children are grouped by age and stimulus duration. Fast corresponds to the mean accuracy of the 100 ms and 150 ms durations. Slow corresponds to the mean accuracy of the 200 ms and 250 ms durations.

| Organism | Accuracy | CI Lower | CI Upper |
|---|---|---|---|
| 3yrold-fast | 0.72 | 0.65 | 0.79 |
| 3yrold-slow | 0.84 | 0.77 | 0.90 |
| 4yrold-fast | 0.74 | 0.65 | 0.82 |
| 4yrold-slow | 0.86 | 0.79 | 0.93 |
| 5yrold-fast | 0.83 | 0.78 | 0.89 |
| 5yrold-slow | 0.93 | 0.89 | 0.97 |
| Adult-fast | 0.99 | 0.98 | 1.00 |
| Adult-slow | 0.98 | 0.94 | 1.00 |
| VoneNet_FF-EcoSet | 0.54 | 0.53 | 0.55 |
| VoneNet_FF-EcoSet-Style | 0.57 | 0.56 | 0.59 |
| VoneNet_R-EcoSet | 0.66 | 0.64 | 0.68 |
| VoneNet_R-EcoSet-Style | 0.67 | 0.65 | 0.69 |
| ResNext50_SAY | 0.60 | 0.59 | 0.62 |
| CVCL | 0.57 | 0.56 | 0.58 |
| ConvNext-ImageNet1k | 0.69 | 0.67 | 0.71 |
| ViT-ImageNet1k | 0.61 | 0.59 | 0.62 |
| ViT-CLIP | 0.99 | 0.98 | 0.99 |
| ResNet50-ImageNet1k | 0.67 | 0.65 | 0.70 |
| ResNet50-ImageNet21k | 0.82 | 0.80 | 0.84 |
| ResNet50-YFCC15m | 0.73 | 0.71 | 0.75 |
| ResNet50-CLIP | 0.91 | 0.89 | 0.92 |

**Supplementary Table 3.** Human and model accuracy on the perturbed condition. CI Lower and Upper refer to upper and lower bounds of the 95% confidence intervals. Children are grouped by age and stimulus duration. Fast corresponds to the mean accuracy of the 100 ms and 150 ms durations. Slow corresponds to the mean accuracy of the 200 ms and 250 ms durations.

| Organism | Accuracy | CI Lower | CI Upper |
|---|---|---|---|
| 3yrold-fast | 0.63 | 0.54 | 0.72 |
| 3yrold-slow | 0.78 | 0.70 | 0.87 |
| 4yrold-fast | 0.80 | 0.74 | 0.85 |
| 4yrold-slow | 0.89 | 0.84 | 0.94 |
| 5yrold-fast | 0.88 | 0.83 | 0.93 |
| 5yrold-slow | 0.91 | 0.87 | 0.95 |
| Adult-fast | 0.95 | 0.90 | 0.99 |
| Adult-slow | 1.00 | 1.00 | 1.00 |
| VoneNet_FF-EcoSet | 0.58 | 0.56 | 0.60 |
| VoneNet_FF-EcoSet-Style | 0.58 | 0.56 | 0.59 |
| VoneNet_R-EcoSet | 0.61 | 0.59 | 0.63 |
| VoneNet_R-EcoSet-Style | 0.64 | 0.62 | 0.66 |
| ResNext50_SAY | 0.64 | 0.62 | 0.66 |
| CVCL | 0.57 | 0.56 | 0.59 |
| ConvNext-ImageNet1k | 0.59 | 0.58 | 0.61 |
| ViT-ImageNet1k | 0.60 | 0.58 | 0.61 |
| ViT-CLIP | 0.97 | 0.96 | 0.98 |
| ResNet50-ImageNet1k | 0.67 | 0.65 | 0.69 |
| ResNet50-ImageNet21k | 0.82 | 0.80 | 0.84 |
| ResNet50-YFCC15m | 0.66 | 0.65 | 0.68 |
| ResNet50-CLIP | 0.83 | 0.81 | 0.85 |

**Supplementary Table 4.** Human and model accuracy on the deleted condition. CI Lower and Upper refer to upper and lower bounds of the 95% confidence intervals. Children are grouped by age and stimulus duration. Fast corresponds to the mean accuracy of the 100 ms and 150 ms durations. Slow corresponds to the mean accuracy of the 200 ms and 250 ms durations.

| Organism | Accuracy | CI Lower | CI Upper |
|---|---|---|---|
| 3yrold-fast | 0.54 | 0.47 | 0.62 |
| 3yrold-slow | 0.58 | 0.52 | 0.64 |
| 4yrold-fast | 0.72 | 0.66 | 0.78 |
| 4yrold-slow | 0.83 | 0.76 | 0.89 |
| 5yrold-fast | 0.67 | 0.59 | 0.75 |
| 5yrold-slow | 0.78 | 0.71 | 0.85 |
| Adult-fast | 0.90 | 0.85 | 0.95 |
| Adult-slow | 0.97 | 0.93 | 1.00 |
| VoneNet_FF-EcoSet | 0.57 | 0.55 | 0.58 |
| VoneNet_FF-EcoSet-Style | 0.54 | 0.53 | 0.56 |
| VoneNet_R-EcoSet | 0.60 | 0.58 | 0.63 |
| VoneNet_R-EcoSet-Style | 0.66 | 0.64 | 0.68 |
| ResNext50_SAY | 0.60 | 0.59 | 0.62 |
| CVCL | 0.56 | 0.55 | 0.58 |
| ConvNext-ImageNet1k | 0.61 | 0.59 | 0.62 |
| ViT-ImageNet1k | 0.58 | 0.57 | 0.60 |
| ViT-CLIP | 0.83 | 0.81 | 0.85 |
| ResNet50-ImageNet1k | 0.58 | 0.56 | 0.61 |
| ResNet50-ImageNet21k | 0.70 | 0.68 | 0.72 |
| ResNet50-YFCC15m | 0.69 | 0.67 | 0.71 |
| ResNet50-CLIP | 0.74 | 0.72 | 0.76 |

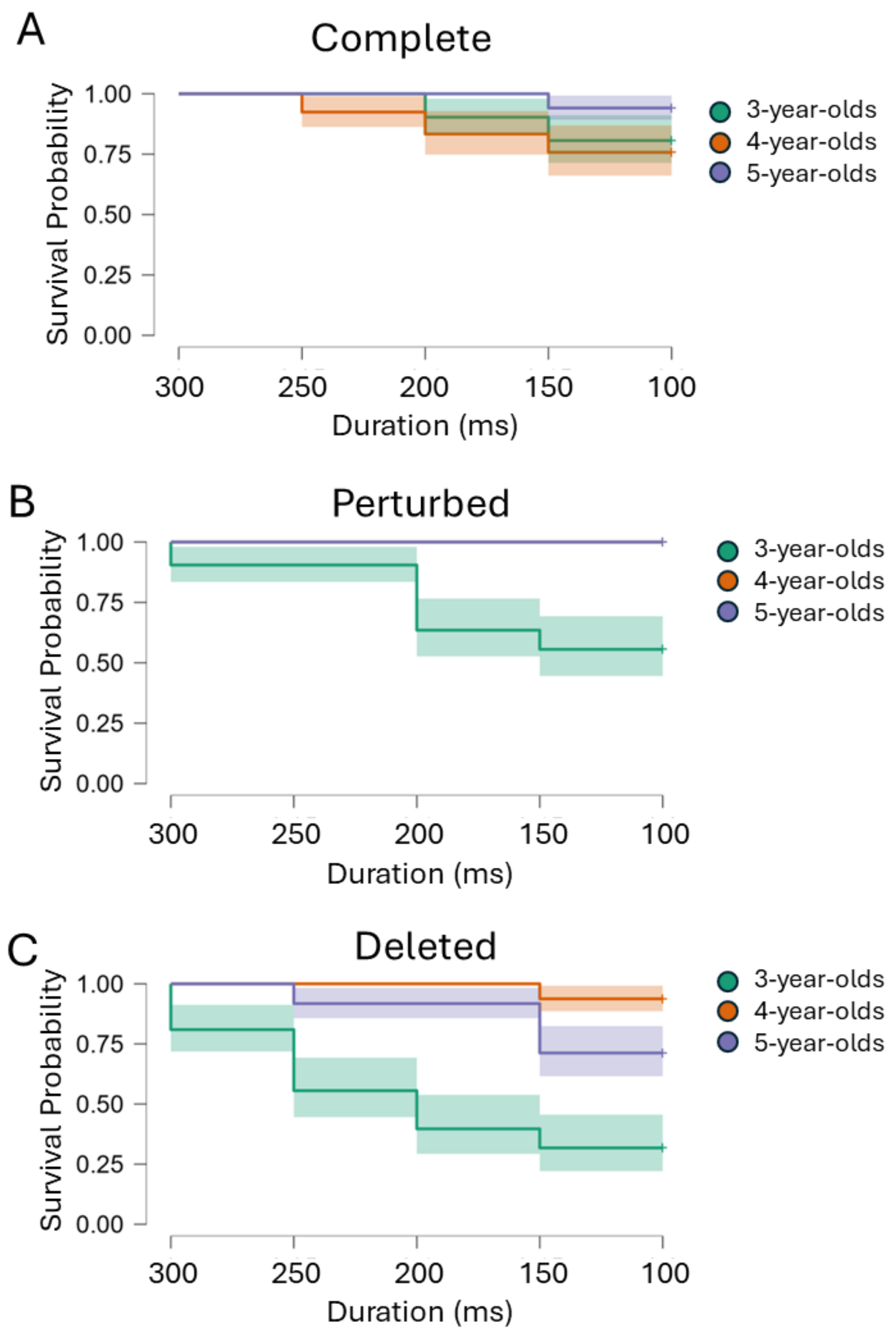


**Supplemental Figure 1.** Survival analyses for (A) complete, (B) perturbed, and (C) deleted conditions for each age group. The X-axis shows the stimulus duration. The Y-axis illustrates the probability of each group reaching each stimulus duration. Note that 4-year-old survival curves are not visible in graph B because they fully overlap with those of 5-year-olds.

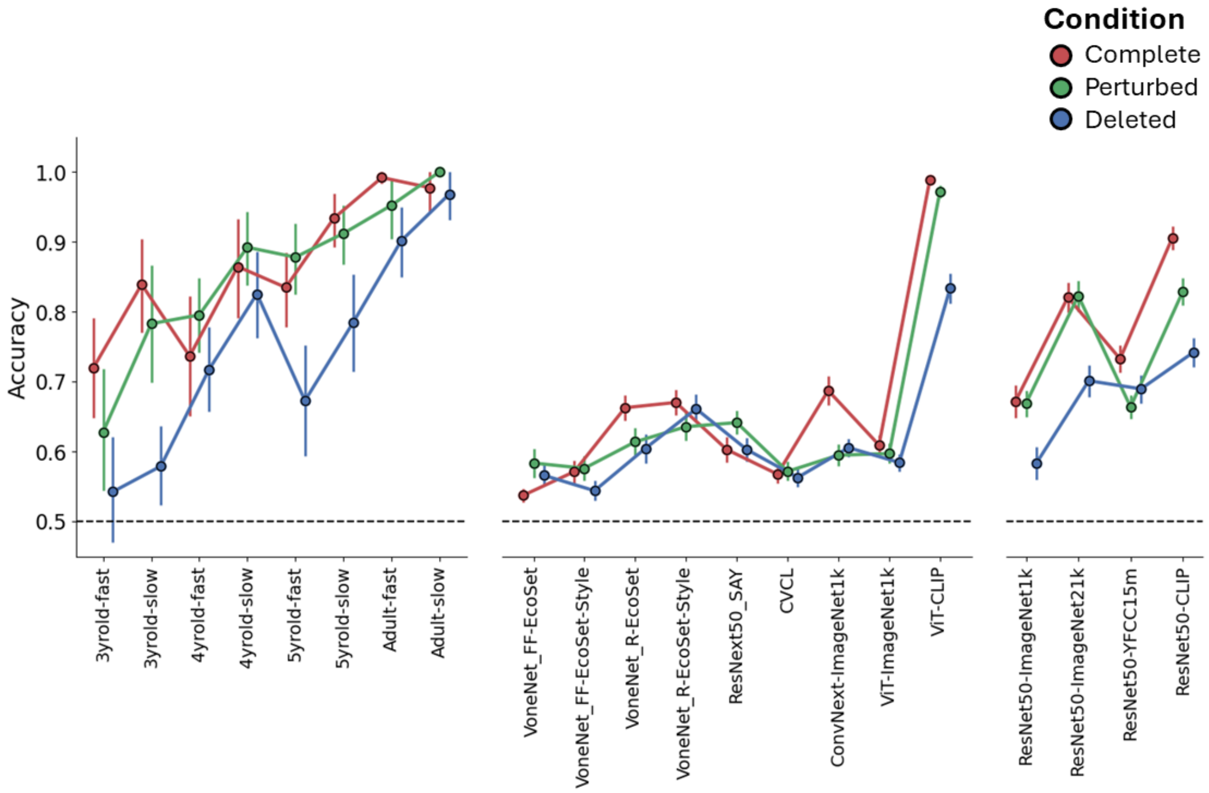


**Supplemental Figure 2.** Performance of humans and models in the (red) complete, (green) perturbed, and (blue) deleted contour conditions. Human data for each age were aggregated into fast (100 ms & 150 ms) and slow (200 ms & 250 ms) stimulus durations. (Left) Humans were compared to (middle) biologically plausible and performance-optimized models, as well as (right) models selected to disambiguate between the contributions of training scale and a vision-language objective. The Y-axis indicates classification accuracy. The dotted line indicates chance performance (0.5) Error bars depict 95% confidence intervals.
